# Transformer Model Generated Bacteriophage Genomes are Compositionally Distinct from Natural Sequences

**DOI:** 10.1101/2024.03.19.585716

**Authors:** Jeremy Ratcliff

## Abstract

Novel applications of language models in genomics promise to have a large impact on the field. The megaDNA model is the first publicly available generative model for creating synthetic viral genomes. To evaluate megaDNA’s ability to recapitulate the nonrandom genome composition of viruses and assess whether synthetic genomes can be algorithmically detected, compositional metrics for 4,969 natural bacteriophage genomes and 1,002 *de novo* synthetic bacteriophage genomes were compared. Transformer-generated sequences had varied but realistic genome lengths and 58% were classified as viral by geNomad. However, the sequences demonstrated consistent differences in various compositional metrics when compared to natural bacteriophage genomes by rank-sum tests and principal component analysis. A simple neural network trained to detect transformer-generated sequences on global compositional metrics alone displayed a median sensitivity of 93.0% and specificity of 97.9% (n = 12 independent models). Overall, these results demonstrate that megaDNA does not yet generate bacteriophage genomes with realistic compositional biases and that genome composition is a reliable method for detecting sequences generated by this model. While the results are specific to the megaDNA model, the evaluate framework described here could be applied to any generative model for genomic sequences.

## Introduction

The widespread development and application of language models has expanded their impact beyond traditional application in natural language processing (NLP). Transformer models, which leverage selfattention mechanisms and deep neural networks, are able to learn the intricate patterns and implicit rules of language. Genomics is essentially a form of language, with nucleotide (and protein) sequences encoding biological information based on the content and order of characters. Transformer-based large language models have proven particularly adept at identifying ‘missing’ annotations for nucleotide sequence data, including for promoters, splice sites, enhancers, chromatin accessibility, transcription factor binding sites, and more (Consens et al., 2023; Ji et al., 2021; Zhou et al., 2023; Dalla-Torre et al., 2023; Avsec et al., 2021). Less common, however, has been the application of transformer-based models for generating novel nucleotide sequences.

In December 2023, Shao published the first version of their language model “megaDNA” for the generation of *de novo* synthetic bacteriophage genomes (Shao, 2023). The megaDNA model leverages the architecture underlying the MEGABYTE model from Meta AI, which is specifically designed for long input sequences (Yu et al., 2023). In the preprint, Shao used a random four bp primer to generate 1,024 sequences longer than 1 kilobase (kb) and demonstrated that a portion of these sequences can be classified as viruses by an AI/ML-based classification framework, show realistic gene lengths and distributions, and encode potentially functional 5’ untranslated regions (UTRs), genes, and promoters.

Genome composition is the reduction of nucleotide sequences into quantitative descriptions of the frequency of specific patterns. These measures can be used to assess the nonrandom distribution of nucleotides in a genome, including measures such as GC content, di- and trinucleotide odds ratios, codon pair bias, and secondary structure elements, among others. RNA viruses have been extensively studied for compositional biases, showing family- and host-dependent patterns (Gaunt and Digard, 2022; Simmonds et al., 2013). While less thoroughly studied, bacteriophages have also shown diversity in GC content and select dinucleotide ratios (Forni et al., 2024). These biases, manifesting in genome-level measurements, were hypothesized to be an appropriate benchmark for megaDNA’s ability to recapitulate higher-order relationships. Identification of a computational method to discriminate transformer-generated from natural sequences would have broad applicability to synthetic biology, biosecurity, and future applications of AI/ML to genomics.

Here, compositional biases in sequences generated by megaDNA were compared to those of natural bacteriophage sequences from the NCBI RefSeq database. Using a variety of approaches, these sequences are shown to be compositionally distinct from the natural sequences upon which they were trained. A variety of methods of varying complexity, from single metric rank-sum tests to 19-feature neural networks, differentiate the two populations with high accuracy on the basis of quantitative compositional metrics alone.

## Results

### Transformer-generated sequence composition is generally dissimilar to that of natural sequences

To compare the composition of natural and transformer-generated bacteriophage genomes, 4,969 RefSeq bacteriophage genomes were downloaded from NCBI and 1,095 synthetic genomes were produced using the megaDNA model with a random four base pair (bp) primer. These synthetic sequences are referred to as “transformer-generated”. Consistent with Shao (2023), analyses were limited to the 1,002 transformer-generated sequences that were greater than 1 kilobase (kb) in length. In contrast to Shao (2023), however, natural sequences with lengths greater than 96 kb were not removed. These longer sequences (n = 721) represent 14.4% of the natural composition dataset.

Natural sequences had a median length of 46 kb, with noted peaks around 6 kb, 40 kb, and 170 kb (Figure 1A). Transformer-generated sequences were on average shorter (median length of 23.7 kb) and displayed a much smoother probability distribution. Length distributions were significantly different (p < 2.2e-308, twotailed Kolmogorov-Smirnov test). The smooth distribution of the transformer-generated sequences is not unexpected, as the predictive approach of megaDNA can infer that the random 4bp primer sequence is located at any position in a theoretical genome. Thus, the sequence output can essentially be thought of as ‘fragments’ that always extend to the 3’ end of the genome. geNomad, an AI/ML-enabled taxonomic classification framework, was used to assess the ability of megaDNA to produce sequences classified as viral by non-kmer approaches (Camargo et al., 2023). As expected, transformer-generated sequences produced significantly lower virus scores than the natural sequences (Figure 1B; median values of 0.72 and 0.98, respectively, p < 2.2e-308, two-tailed MannWhitney U Test). Only 22 transformer-generated sequences had predicted taxonomy, all of which were *Caudoviricetes*. Based on genomad virus scores, transformer-generated sequences were split into low (virus score < 0.7; n = 418), medium (virus score ≥ 0.7 and < 0.8; n = 281), and high quality (virus score ≥ 0.8; n = 303). In this study, geNomad was executed with the`–relaxed` quality parameter to ensure scores were reported for all sequences. This explains the on average higher geNomad score results reported in Shao (2023) where geNomad was almost certainly executed with default quality parameters. The cause of the much lower classification efficiency in this study compared to Shao (2023) is unclear, but could be due to using different versions of geNomad or MMSeqs2. The version numbers were not specified in Shao (2023).

**Figure 1.**
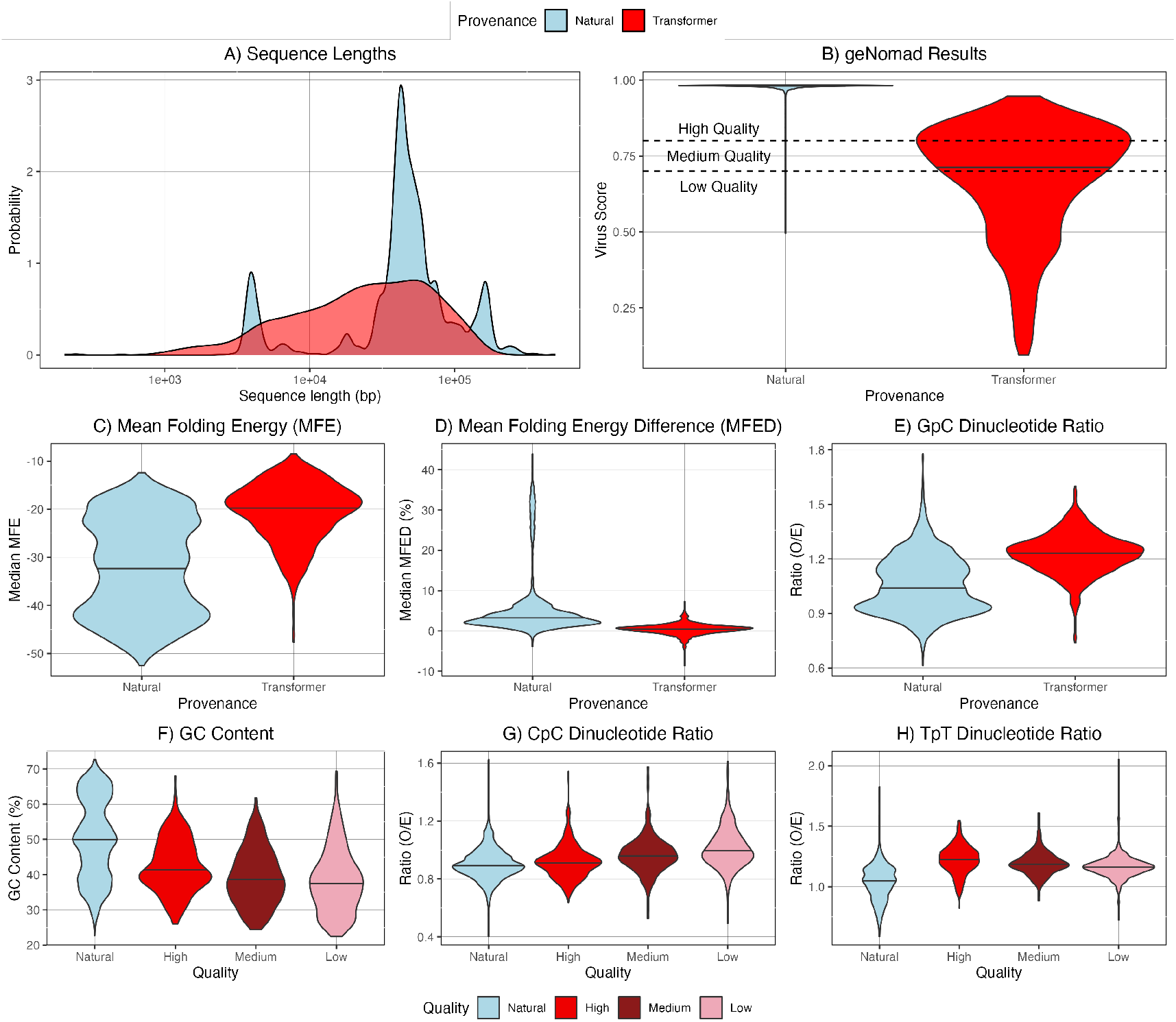
Characteristics of Natural and Transformer-Generated Sequences. For all violin plots, black horizontal bar displays median value. (**A**) Sequence length for natural sequences and all transformer-generated sequences *≥* 1000 bp. Probability distributions significantly different (p < 2.2e-308, two-tailed Kolmogorov-Smirnov test). (**B**) geNomad virus score results. Quality cutoffs of 0.7 and 0.8 are based on parameters described within the software documentation. (**C, D, E**) Differences between select metrics; all distributions significantly different by two-tailed Mann-Whitney U Test (p < 0.0026). (**F, G, H**) Differences between select metrics with transformer-generated sequences grouped according to geNomad virus quality; GC content and TpT Ratio significantly different between natural and high quality sequences (p < 0.0026).

Nineteen compositional metrics were analyzed. These included GC content, odds ratios for all 16 dinucleotides, median mean folding energy (MFE) values for overlapping 120bp subsequences, and mean folding energy difference (MFED) values calculated against dinucleotide shuffled controls. Of the 19 compositional metrics considered, the distributions for natural versus transformer-generated were significantly different for 18 of the metrics after a Bonferroni correction (examples in Figure 1C-E; Figure S1). For dinucleotide odds ratios, absolute differences in medians ranged from 0.0003 (ApA) to 0.194 (GpC). Transformer-generated sequences were consistently more GC-poor (median values of 39.2% versus 49.9%), had lower folding energy (median MFE values of -19.7 versus -32.2), and had MFED values much closer to the null expectation (0.005 versus 0.0324) than natural sequences. Differences in MFED values were particularly stark, as only 3% of transformer-generated sequences displayed MFED values as high as the median natural sequence (Figure 1D).

The neural network classification module in geNomad uses 4-mer encoded subsequences, which may lead to scores reflecting compositional differences. Supporting this, 72% of comparisons of composition metrics between low, medium, and high quality transformer-generated sequences were significantly different (Table S1). If the geNomad scores were implicitly detecting composition bias towards natural sequences, it would be expected that high quality sequences would be most similar to natural, low quality would be the most different, and medium quality would be somewhere in the middle (as in Figure 1F). Indeed, this pattern was observed 36.8% of the time, significantly more frequently than would be expected by chance (p = 7.6e-5, binomial distribution). However, these transformer-generated sequences still had significantly different composition metric distributions than natural sequences for 17 of the 19 measured metrics (Figure 1F-H; Table S1).

Overall, the transformer-generated sequences display significantly altered composition metrics when compared to the global bacteriophage population. Additionally, a portion of the sequences are classified as viral by geNomad and there is some evidence of a relationship between virus score and ‘natural-like’ composition.

### Bacteriophage sequences cluster based on provenance and taxonomic relationships

It is generally observed that genome composition varies between viral families (Simmonds et al., 2013). Without knowledge as to the specific sequences used to train megaDNA, it is possible that the training sequences were heavily biased toward an individual family and, as such, comparisons against a population of bacteriophage sequences that does not reflect the same bias - as done in the previous section - are an unfair and harsh comparison. To evaluate whether transformer-generated sequences are simply many members of a specific viral family to which the model is overfitting, principal component analysis (PCA) of the compositional metrics was performed. PCA was preferred to the use of raw metrics as the nature of dinucleotide ratios leads to many correlations between them (Figure S2).

PCA was performed using data for all sequences. For cluster and distance analysis and visualization, however, the comparisons were limited to only transformer-generated sequences and those bacteriophage genomes whose GenBank file specified the viral family (n = 2080) (Figure 2A). To determine an individual point’s location in the 19-dimensional space returned by the PCA, the raw PC values were weighted according to the square of their respective eigenvalues (colloquially, the percent of variance explained by that PC). To evaluate the extent of clustering for the 28 families with ≥ 10 members in the dataset (including transformer-generated as a family), the frequency with which a data point’s nearest neighbor in the weighted 19-dimensional space was a member of the same family was compared to expectations from random associations. For all families, individual members were more likely to be ‘close’ to other members of the same family than random sequences in the dataset (Table S2). While all significant by binomial distributions (p < 0.002), the strength of clustering varied between families. For *Blumeviridae*, only five of 34 members clustered. For *Demerecviridae*, all 84 members of the family clustered with another member of that family, likely due to a combination of the family’s low CpG and high TpA ratios compared to all other sequences (0.82 vs 1.00 and 0.96 vs 0.77, respectively, p = 6.89e-20 and p = 1.99e41, two-tailed Mann-Whitney U Tests). Transformer-generated sequences clustered at a rate of 95.2% (n = 954/1002). This result was also demonstrated in a PCA that only included dinucleotide ratios and GC content (Figure S3). There were no differences in cluster rate between low, medium, and high quality transformer-generated sequences (93.1%, 96.4%, and 97.0%, respectively). These results confirmed that PCA extracted family-specific compositional traits from the natural sequences.

**Figure 2.**
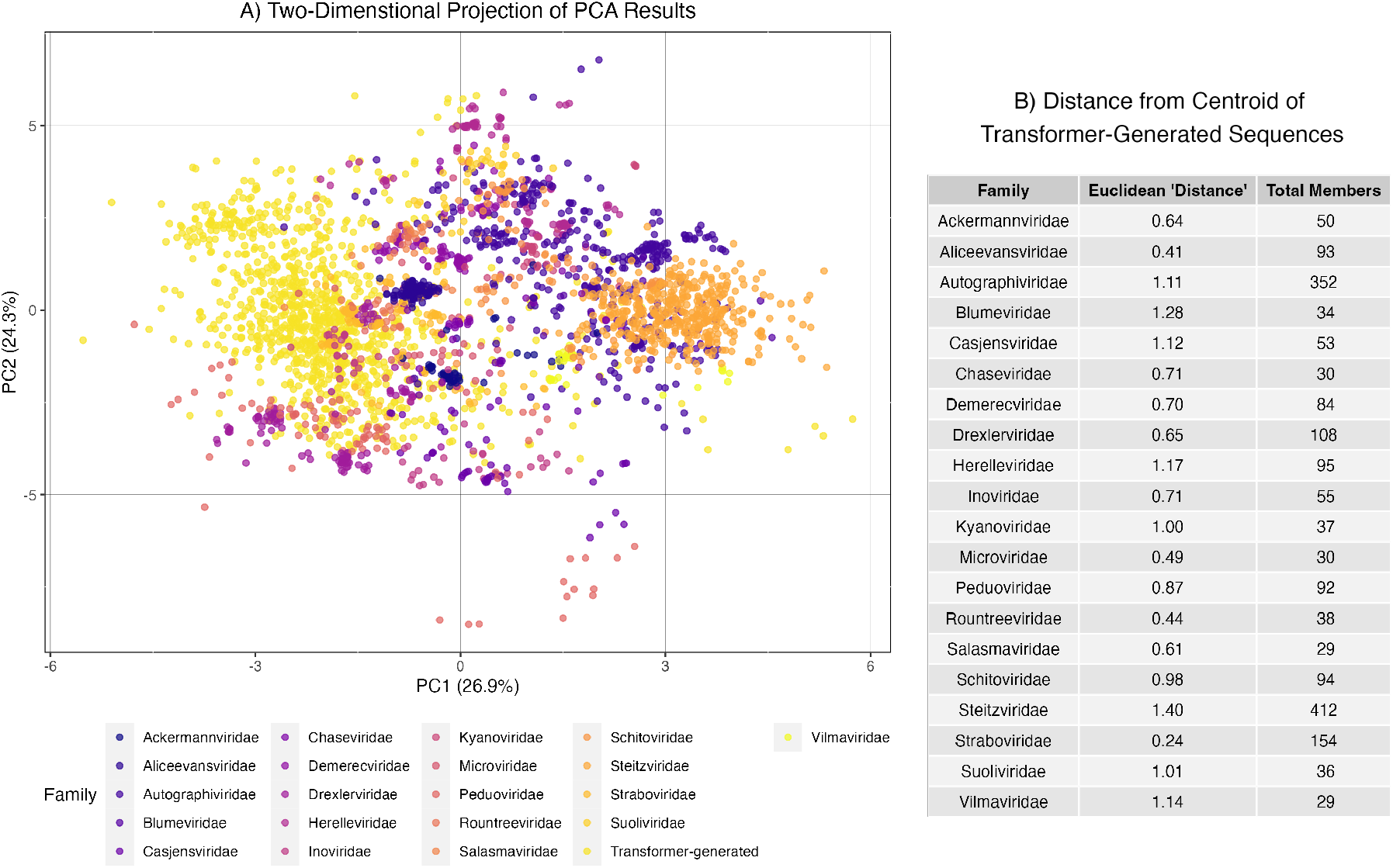
Principal Component Analysis (PCA) of Compositional Metrics. (**A**) Two-dimensional projections of PCs 1 and 2 limited to sequences from families with ≥ 25 members, colored by taxonomy. The values for this projection have **not** been weighted according to the square of the eigenvalues. Percentages in x-axis and y-axis titles indicate percentage of variance explaned by given PC. Kindly note that the two-dimensional projection can be misleading as to the true location of a data point in the 19-dimensional space created by the PCA. (**B**) Unitless weighted Euclidean distance measures from centroid of family to centroid of transformer-generated sequences.

To assess to which viral family transformer-generated sequences were most similar, a weighted Euclidean distance was calculated between centroids of each viral family that contained ≥ 25 members. The unitless distance measures ranged from 0.24 (*Straboviridae*) to 1.40 (*Steitzviridae*) (Figure 2B). Of important note, the ‘closeness’ of *Straboviridae* and *Demerecviridae* to the transformer-generated sequences in the 19dimensional space is especially surprising as these particular sequences from these families should not be present in the training set used for megaDNA as they are beyond the stated 96 kb size exclusion threshold. Further, there were still significant differences between Straboviridae and transformer-generated sequences for 12 of the 19 metrics (p < 0.0026, two-tailed MannWhitney U Test), strongly suggesting transformer-generated sequences are not simply generating sequences with the same compositional bias as their most similar family.

All together, the PCA results suggest transformer-generated sequences occupy an independent compositional niche. This niche likely reflects the average of the compositional biases of the sequences within the training set and suggests that sequences produced by megaDNA revert to their compositional mean rather than reflect the compositional bias of a single sequence or family in the training set. This hypothesis could be further assessed by priming megaDNA with fragments of sequences of known taxonomy and assessing the degree to which the produced sequence diverges from the ground truth fragment not seen by the model.

### A simple neural network differentiates transformer-generated and natural sequences with very high accuracy

If transformer-generated sequences truly occupied an isolated compositional niche, then they should be able to be identified on the basis of compositional metrics alone. To test this hypothesis, a neural network with a relatively simple architecture of two hidden layers was trained to differentiate natural and transformer-generated sequences based on compositional metrics. Twelve models were generated on an identical 80:20, train:test population using all 19 compositional metrics. These ‘total’ models displayed a median overall accuracy of 97.0% (sensitivity = 93.0%, specificity 97.9%). Importantly, the neural network input dataset reflected the natural vs transformer-generated sequence imbalance seen throughout this study. This asymmetry leads to a ZeroR model accuracy of 82.6%.

To determine if the neural network classification performance was dependent on the number of total metrics or a specific metric potentially limiting its generalizability beyond megaDNA new models were trained on random feature subsets. For each total feature count from 1 to 18, 12 models were trained using a random selection of features (216 models total). Accuracy for these ‘incomplete’ models was assessed using the same test population as for the ‘total’ models. Overall performance was seemingly dependent on the near-full breadth of features, with model performance only reaching 95% of its maximum median accuracy (a threshold of 96.6%) after 16 features were present (Figure 3A).

**Figure 3.**
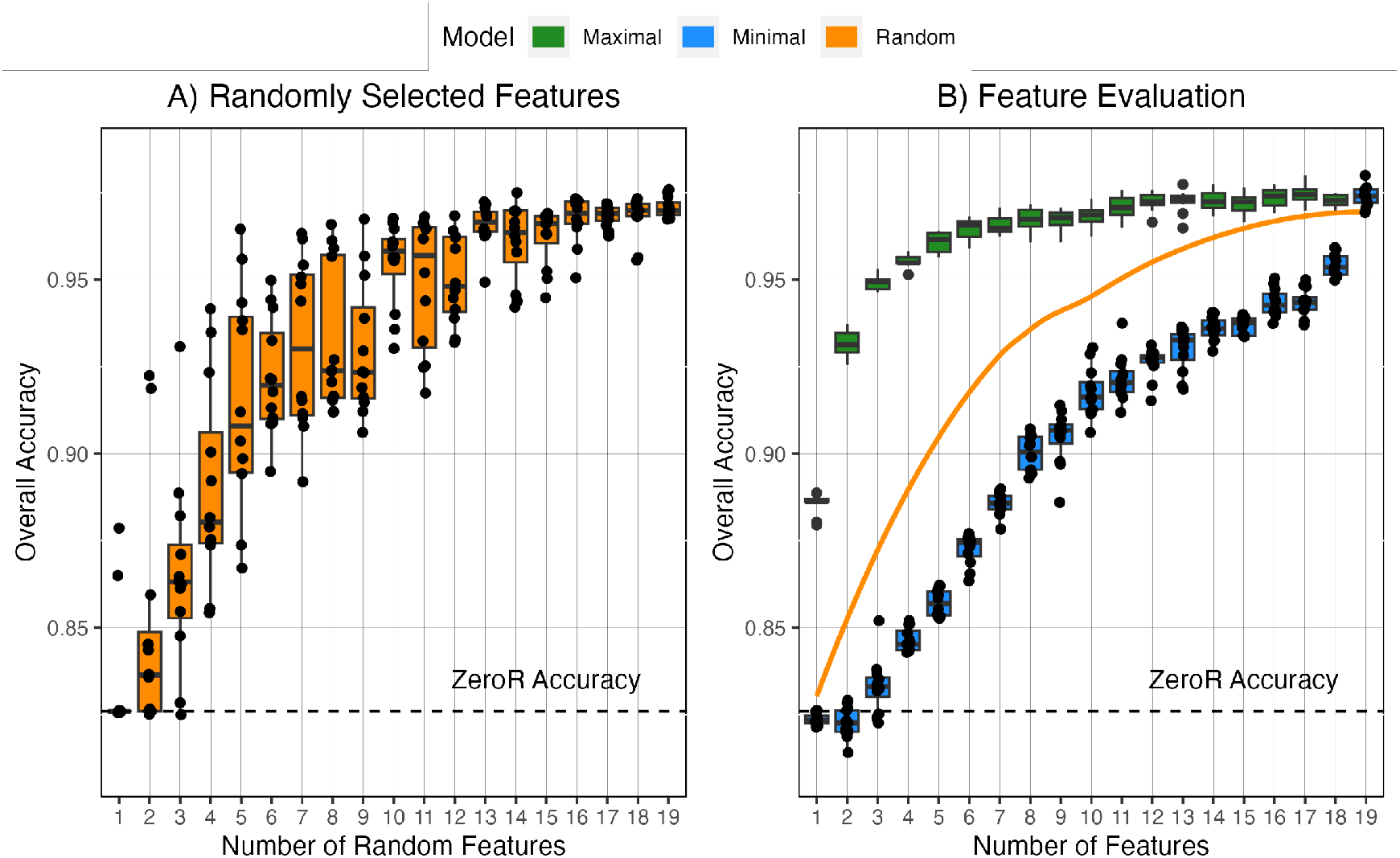
Discrimination of Transformer-Generated Sequences with Neural Networks Trained on Varied Numbers of Features. (**A**) Overall classification accuracy for models built with a random selection of features. Boxplot displays distribution statistics for n = 12 replicates. (**B**) Results from predictive feature evaluation (described in detail in methods). Feature order for maximal and minimal models available in Figure S4. Orange line displays LOESS regression for random features using the R stats package.

There was stark variability in the performance of the ‘incomplete’ models that must be explained by which features were included. For example, a model with only five random features displayed an accuracy level of 96.5%, which outperformed exactly half of the models with 14 random features. To determine which metrics were the most and least predictive, new features were iteratively added to baseline models by testing and selecting the metric which led to the greatest or least increase in accuracy (Figure 3B, Figure S4). Akin to a greedy algorithm, this method selects the locally optimal solution at each step, which may not reflect the ‘true’ global order of feature addition for maximal accuracy.

The results of the predictive feature evaluation were unambiguous as to the importance of MFED in discriminating natural and transformer-generated sequences. Not only was MFED the most predictive single metric (supported by Figure 1D), but its exclusion led to an 18feature model displaying lower median accuracy than that of a model that only included MFED and GpT, CpC, and ApC ratios. Interestingly, for the maximum accuracy feature evaluation, 95% of the improvement from ZeroR to the total model is achieved through the addition of only eight features (compared to 16 for the random selection). Combined with the significant drop in performance of models not including MFED, this result is suggestive that the performance contribution of dinucleotide ratios is saturated quickly, potentially due to the high correlation rates between dinucleotide ratios (Figure S2). Future studies may need to investigate compositional metrics beyond dinucleotide ratios, MFE, and MFED values to achieve higher accuracy discrimination.

## Discussion

Effectively leveraging generative AI to overcome problems in biology could lead to breakthroughs across multiple domains. While generative approaches have seen impact in fields such as drug discovery, protein folding, and designing functional ribozymes (Zeng et al., 2022; Eguchi et al., 2022; Sumi et al., 2024), the use of transformer models specifically is nascent. In a future state, iterative design pipelines could harness the power of transformer models for generation, complemented by encoder models for functional prediction and evaluation. Therefore, establishing methods to rapidly evaluate the outputs of these early models *in silico* and enable refinement of their architecture and training approaches will advance the development of the field.

As generative AI continues to proliferate, there are legitimate concerns over it’s misuse. While much of the commentary has focused on using LLMs for knowledge discovery and synthesis (Patwardhan et al., 2024; Mouton et al., 2023), other authors have identified the risk of language models for accelerating the design of bio-logical weapons (BW) (Sandbrink, 2023). As described by Sandbrick (2023), these ‘biological design tools’ may enable circumvention of known biosecurity measures or BW detection methodologies. While important, discussions around the appropriate oversight of the development of generative AI models for biology are outside the scope of this paper. In general, this author is of the opinion that the risks poised by models that can learn currently unknown functional properties that influence pathogenicity, toxicitity, and transmissibility of viruses is higher than that of transformer models that can generate new sequences from noise. Studies of the former would be akin to the concerns raised by the experiments conducted by Herfst et al. (2012), where determinants of H5N1 influenza virus airborne transmissibility could be inserted into the backbone of an existing influenza virus (Herfst et al., 2012). This conclusion is also partly informed by the specialist knowledge and equipment as well as large amount of capital required to synthesize and evaluate a transformer-generated virus in a laboratory setting.

While the notion of transformer-generated pathogens is somewhat fantastical (and the analysis in this paper demonstrates those capabilities are not available in the near term), there are minor risks of the sequences themselves being used maliciously. For example, fully synthetic sequences could be used to intentionally poison public sequence databases. It is therefore pertinent to be able to discern transformer-generated sequences those which have been manufactured through human intervention from those arising from natural processes. Similar to the large number of tools available to detect ChatGPT generated text (Højris Bæk, 2023), the development of methods that can discern ‘fingerprints’ of transformer-models in sequence data should be prioritized.

This paper has demonstrated that these markers exist, at least in the circumstances where labelled data is available. The megaDNA generated sequences are clearly compositionally distinct from natural bacteriophage genomes, with this finding being consistent regardless of the tested metric (dinucleotide ratios, GC content, secondary structure). Overall, the findings point to two classes of compositional shortcomings in the megaDNA outputs:

1) The distinctness of the megaDNA dinucleotide ratios and GC content (calculated across whole sequences) are suggestive that the model overgeneralizes the distinct compositional biases of its training sequences (Figure S3). Rather than generating sequences closely resembling specific families represented in the training set, the model’s outputs appear to be a weighted average of all experienced compositional pressures of its training sequences. It’s possible, albeit unlikely, that this issue can be overcome with additional training data. It does, however, suggest that a path forward for the generation of whole genomes that resemble members of existing virus families may require the development of family-specific models. This may also be achieved by prepending taxonomic labels to sequences in the training dataset, as done elsewhere (Nguyen et al., 2024). However, the low data availability for the vast majority of viral families will challenge their development in the short-term.

2) The dramatic difference in MFED values strongly supports a failure of megaDNA to learn associations between neighboring bases that govern secondary structure (and other non-random base distributions). This issue may arise due to MEGABYTE’s impressive long-context capabilities that allow it to reconstruct the spatial organization of a bacteriophage genome (e.g., promoters and genes, as seen in Shao (2023)) also result in it paying less attention to the finer, more localized interactions that underlie these relationships. Resolving this particular issue may require migrating to a different architecture than MEGABYTE.

While this manuscript was in its final preparations, Zhao et al. (2024) preprinted a manuscript describing GenerRNA, their transformer-based model for the generation of synthetic RNA (Zhao et al., 2024). Based on a configuration similar to the OpenAI GPT2 architecture, the authors demonstrate that their approach generates short RNAs with realistic secondary structure profiles and protein-binding characteristics. A rapid analysis using natural and generated sequences deposited to their GitHub repository (https://github.com/pfnet-research/GenerRNA) demonstrated that GenerRNA was substantially more similar to its training data than megaDNA sequences with only two composition metrics having significant differences (S6). Importantly, while GenerRNA sequences did display significantly lower MFED values than their training sequences (as seen for megaDNA), the magnitude of difference was minor (0.070 versus 0.078). The raw MFED value being much greater than 0 is highly suggestive that, unlike megaDNA, the architecture used by GenerRNA has successfully ‘learned’ to maintain the local sequence relationships that underpin RNA secondary structure. This result is suggestive that token-based approaches, which may not be at the single nucleotide resolution, are not inherently flawed at maintaining local context features. Lastly, while the authors do not clarify the token limit of GenerRNA, the 1024 token limit of GPT-2 would be unlikely to recapitulate the genome organization that megaDNA excelled at, so future applications of generating whole genomes will need to balance the pros and cons of each architecture or explore novel methods for combining the two approaches.

The intent of this research was to perform a fair and unbiased assessment of the megaDNA model. While the results focus on potential shortcomings of the current version of the model, its remarkable success in other aspects such as generating potentially functional promoters and maintaining realistic genome organization should be celebrated. Bin Shao, the creator of megaDNA, should also be recognized for his willingness to open-source an in-development model.

The findings presented herein are specific to the megaDNA model weights retrieved in late December 2023 and are unlikely to generalize broadly. For future research, the methods and approach described here can be used as a general framework to assess the ability of generative AI to recapitulate the compositional biases inherent to RNA and DNA sequences.

## Methods

### Creation of Transformer-Generated Sequences

Transformer-generated sequences were produced using the model and model inference codes from Shao (2023) as of Dec 27, 2023: https://github.com/lingxusb/megaDNA. The inference code was modified to encapsulate the `model.generate()` call within a `with torch.no_grad()` statement to reduce memory consumption. Model inference was performed on an NVIDIA V100.

### Identification of Natural Bacteriophage Genomes

Shao (2023) does not list the accession numbers for all sequences used to train the transformer model described in the preprint. To ensure only high-confidence and high-quality sequences of real bacteriophages were used for comparison, n = 4,984 RefSeq sequences containing “phage” within the organism’s name were identified from the NCBI virus database in December 2023. There is a high likelihood these sequences are present within the training set used by Shao (2023). GenBank files for each accession number were downloaded from NCBI using the entrez API available via Biopython (Cock et al., 2009).

### Sequence Compositional Analysis

Sequence composition analysis was performed using custom python scripts. GC content and dinucleotide odds ratios were calculated for whole genomes. Structural metrics (Minimum Free Energy (MFE) and Minimum Free Energy Difference (MFED)) were calculated for overlapping subsequences for each genome from which median values were extracted. Briefly, a sequence was exploded into overlapping k-mers of prespecified length and step size and stored in a numpy array (Harris et al., 2020). Metrics were calculated for each k-mer and values assigned to the index of the midpoint nucleotide of the k-mer in the original sequence. For this study, a k-mer size of 120bp and step size of 10bp were used.

#### GC Content

GC content was calculated as the sum of guanine and cytosine bases in the sequence divided by the length of the sequence.

#### Dinucleotide Odds Ratios

Dinucleotide odds ratios were calculated based on the expectation of random assortment of mononucleotides of fixed frequencies. Therefore, ratios of observed versus expected dinucleotide frequencies can be derived for any sequence with a known mononucleotide frequency. Odds ratios less than one represent suppression while ratios greater than one indicate overexpression.

Formally, the dinucleotide ratio for any two nucleotides X and Y can be calculated as:

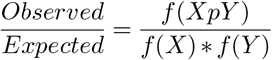

Where *f* (*XpY*) is the observed frequency of dinucleotide XpY, *f* (*X*) is the observed mononucleotide frequency for mononucleotide X, and *f* (*Y*) is the observed mononucleotide frequency of Y.

#### Minimum Free Energy (MFE)

MFE values were calculated using the `RNA.fold()` function from the Vienna RNA python package (Lorenz et al., 2011). While designed for single-stranded (ss) RNA, these algorithms were applied equally to ss and double-stranded (ds) RNA and DNA sequences. See the next section for a disclaimer about deriving structural metrics for dsDNA genomes.

#### Minimum Free Energy Difference (MFED)

MFED values were calculated as the percentage difference between the MFE of a sequence and those of a selection of permuted controls, as performed elsewhere (Simmonds et al., 2004). There are multiple approaches for creating permuted controls that can lead to inconsistent conclusions (Clote et al., 2005). For every 120bp oligomer, a dinucleotide shuffling algorithm was used to generate a set of 105 permuted controls against which the MFE of the oligomer was compared (see Supplementary Methods).

Of note, Shao describes 98% of the sequences used in the model training dataset as being members of the *Caudovirales* class of dsDNA phages. As these are expected to be fully base-paired, it is reasonable to question the use of MFE or MFED values to describe their composition. However, the natural genome sequences used in this study, when treated as ssRNA, were consistently more structured than would be expected by chance (97.6% with median values > 0 (n = 4850/4969); p < 2.2e-308, binomial distribution). This result confirms that MFE and MFED results can be used as measurements of the non-randomness of bacteriophage base composition even if their contribution to nucleic acid binding dynamics is unclear.

#### Dinucleotide Shuffled Sequence Generation

Dinucleotide shuffling is a method of shuffling sequences in a manner that conserves the non-random distribution of dinucleotide frequencies of the input sequences. Here, dinucleotide shuffling was conducted using an open-source implementation of the altschul-erikson algorithm written in python (Altschul and Erickson, 1985): https://github.com/wassermanlab/BiasAway/.

### Virus Score Prediction

All sequences were processed using geNomad v1.7.4 with the `– relaxed` parameter and the MMseqs2 release 15-6f452 (Camargo et al., 2023; Steinegger and Söding, 2017).

### Principal Components Analysis (PCA)

PCA was conducted on scaled compositional metrics using the `prcomp` function in R version 4.2.0 (R Core Team, 2021). Group centroids were calculated as the mean value of each PC and expressed as a matrix of coordinates. The ‘distance’ between points or group centroids in the 19-dimensional space was calculated as a Euclidean distance measure weighed based on the square of each principal component’s (PC) eigenvalue. Weighed Euclidean distance was used in place of downselecting PCs using the Kaiser criterion or a scree plot due to criticism of the two methods being arbitrary or subjective (Zwick and Velicer, 1986; Fabrigar et al., 1999). Further, Euclidean distance was preferred to Mahalanobis distance as the raw data was scaled to unit variance before the PCA was performed. As the square of the eigenvalue is equivalent to the proportion of variance explained by each PC, distance measures are dominated by the PCs that explain the greatest proportion of the total variance.

Nearest neighbors for individual points were identified using function `get.knn` from the `FNN` package. The null hypothesis of random association predicts that the frequency of a nearest neighbor being from the same family of a given point is equal to:

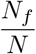

Where *N*_*f*_ is the total number of points for family *f* and *N* is the total number of points in the dataset.

### Predictive Neural Network

To create a predictive neural network in python, compositional metric data was loaded into a pandas DataFrame and all features were scaled using using `StandardScaler()` from `scikit-learn`(Pedregosa et al., 2011). Sequence provenance (Natural versus Transformer) was encoded as 0 or 1, respectively, and an 80:20 training:test split was used. The neural network architecture used the `Sequential()` model from the TensorFlow Keras API (Martín Abadi et al., 2015). The model consisted of two hidden layers with 32 and 16 neurons, respectively, both with rectified linear unit (ReLU) activation functions, and an output layer with one neuron and a sigmoid activation function. The model was compiled using binary crossentropy loss and the Adam optimizer and trained using 10 epochs and a batch size of 32. Model performance on the testing data was expressed as total accuracy, sensitivity for transformer-generated sequences, and specificity for natural sequences.

### Predictive Feature Selection

Predictive feature evaluation was performed iteratively to maximize (or minimize) the predictive capability of models with limited features. This iterative approach began with a baseline model containing no features. During each step, each of the remaining features was added to the existing set of features and used to train 12 independent models. Following the evaluation of each feature, the metric whose addition resulted in the highest (or lowest) average increase in accuracy was selected and incorporated into the model. This partial model was used to continue the iterative process until all features had been selected.

### Statistical Analysis

Between group comparisons were made using twotailed Mann-Whitney Tests. Binomial distributions were used in instances where binary outcomes could be derived (e.g., k-nearest neighbor analysis of family clustering, order of relationships within geNomad, etc.).

## Acknowledgements

My utmost gratitude is extended to Bin Shao for preprinting the article upon which this work is based and for open-sourcing the sequence generation model. I thank Vijay Narayan for assistance with the sequence processing scripts and acknowledge Steve Royle, Ricardo Henrique, and Theo Sanderson for this preprint template (available at: https://github.com/quantixed/manuscript-templates). ChatGPT assisted with code generation and debugging. Portions of this work were supported by internal research funding provided by the the Johns Hopkins University Applied Physics Laboratory.

## Supplementary Methods

### MFED sample size optimization

MFED values are essentially an estimate of the relative ranking of a given MFE value within a pool of MFE values derived from ‘similar’ sequences. Dinucleotide shuffling returns a single ‘similar’ sequence that is the product of a valid solution to the Eulerian paths as described in Altschul and Erickson (1985). The shuffled sequence is therefore a single sample of a large population of potential solutions. As MFED values are derived by comparing an input sequence against a selection of shuffled controls, the extent to which that selection represents the characteristics of the population is key to the reproducibility and accuracy of the MFED estimate.

The number of shuffling solutions grows exponentially depending on input sequence length (Figure S5A), and can therefore effectively be treated as infinite for the sequence lengths relevant to this study. Based on the Central Limit Theorem, increased sample sizes will reduce the variability of an MFED estimate compared to other estimates drawn from the same population. However, larger sample sizes also incur higher computational costs and thus a balanced sample size must be selected. To measure the relationship between sample size and MFED estimate variability and identify an optimal sample size, random sequences of length *l* ranging from 100bp to 400bp were generated and MFED values were processed with varying sample sizes. Values of *l* included 100bp, 150bp, 200bp, 275bp, and 400bp and 15 replicates were generated for each length.

For each sequence, 15 MFED values were derived based on a shuffling sample size of *i*. Values of *i* ranged from 10 to 100 with step sizes of 10. From this population of 15 distinct MFED estimates, the standard deviation *s* was measured. This process was repeated for 15 total *s* estimates. The median *s* value for each value of *i* was carried forward for further analysis, creating a two column dataframe of *i* versus median *s*. The *s* values were normalized based on the maximum median *s* value and a linear regression of *s* vs *i* was performed. While likely not optimal, a linear equation was used to ensure a monotonically decreasing relationship. The resultant equation describes the number of samples *i* necessary to achieve a reduction of Y% compared to the maximum *s* value for that sequence. For every integer value of Y from 1 to 99, the median estimate of *i* for all input sequences of length *l* was collected.

### GenerRNA Evaluation

This evaluation incorporated the 2000 natural and 2000 generated sequences listed within `MFE_distribution_Fig4a.csv` in the GenerRNA GitHub repository (https://github.com/pfnet-research/GenerRNA) as of commit ab7f470 on January 25, 2024. These were subsetted to only those sequences without ambiguous nucleotide codes, yielding 1,939 natural and 1,942 generated sequences for the analysis. GC content, dinucleotide odds ratios, MFE, and MFED values were calculated for whole sequences (rather than 120bp subsequences for MFE and MFED as done for megaDNA). This approach was selected to be similar to that done in the GenerRNA preprint. MFED was calculated with only 20 iterations to save computational time given size of input sequences. PCA and cluster analysis were performed as in methods.

## Supplementary Results

### MFED sample size optimization

As expected, the relationship between sample size and MFED estimate variability was logarthmic, with progressively diminishing impacts on variability for increasing sample sizes (Figure S5B). Results were consistent across all sequence lengths tested. From these results, a shuffling sample size of 105 was chosen to be used in this study as this was shown to reduce the value of *s* by approximately 89% for all sequence lengths. Beyond this level, the diminishing returns of increased sample size were deemed to be an inefficient use of computational resources and time. There is no ‘correct’ sampling size for dinucleotide shuffling and future uses of this method need to balance accuracy with available resources.

### GenerRNA Evaluation

For the 19 compositional metrics analyzed, the distributions of natural versus generated sequences were significantly different for only two after a Bonferroni correction (GpC ratio and MFED, Figure S6). Consistent with the findings of Zhao et al. (2024), there were no differences in distribution of raw MFE values. However, the algorithms used by Zhao et al. (2024) to conclude there were no differences in structure are inadequate compared to dinucleotide shuffling (see Clote et al. (2005)). In contrast to the authors’ claims of parity between natural and generated sequences, MFED values of generated sequences were significantly lower than those of natural sequences (0.070 versus 0.078, respectively, p = 0.000062, two-tailed Mann-Whitney U Test). Lastly, generated sequences did not cluster with themselves greater than expected by chance (p = 0.14, binomial distribution; Figure S7). Compared to megaDNA, the composition of GenerRNA model outputs are substantially more compositionally similar to their training data.

**Figure S1.**
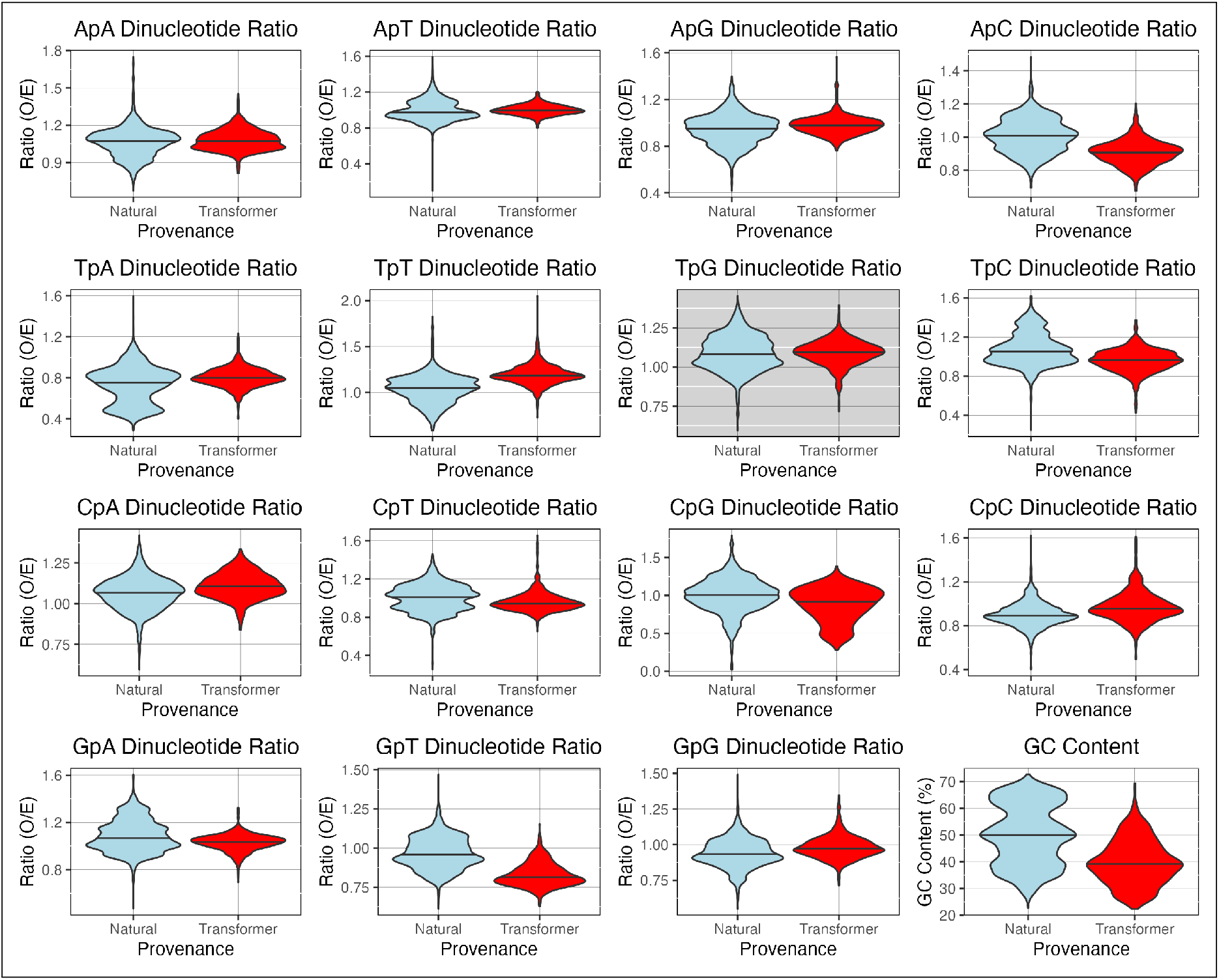
Comparisons of Compositional Metrics between Natural and Transformer-Generated Sequences. Inclusive only of metrics not displayed in Figure 1. All distributions significantly different by two-tailed Mann-Whitney U Test (p < 0.0026) except for TpG dinucleotide ratio (colored grey).

**Table S1.** Distributions of Composition Metrics. Table split into two for ease of viewing. (**B**) Values indicate p value from two-tailed Mann-Whitney U Test. Green highlighted cells indicate comparisons that are statistically significant after Bonferroni correction (p < 0.0026 for Natural vs Transformer or p < 6.6e-4 for all others).

| Metric | A) Median values |  |  |  |  |
| --- | --- | --- | --- | --- | --- |
|  | Natural | Transformer-generated |  |  |  |
|  |  | All | High quality | Medium quality | Low quality |
| GC Content | 0.499 | 0.392 | 0.410 | 0.384 | 0.374 |
| ApA ratio | 1.074 | 1.074 | 1.060 | 1.065 | 1.088 |
| ApG ratio | 0.952 | 0.979 | 0.975 | 0.984 | 0.979 |
| ApT ratio | 0.973 | 0.997 | 1.017 | 0.995 | 0.987 |
| ApC ratio | 1.010 | 0.908 | 0.924 | 0.908 | 0.890 |
| GpA ratio | 1.070 | 1.035 | 1.041 | 1.041 | 1.029 |
| GpG ratio | 0.933 | 0.972 | 0.958 | 0.967 | 0.999 |
| GpT ratio | 0.958 | 0.813 | 0.791 | 0.812 | 0.836 |
| GpC ratio | 1.040 | 1.234 | 1.238 | 1.239 | 1.221 |
| TpA ratio | 0.756 | 0.799 | 0.776 | 0.800 | 0.814 |
| TpG ratio | 1.082 | 1.094 | 1.117 | 1.103 | 1.079 |
| TpT ratio | 1.050 | 1.183 | 1.225 | 1.187 | 1.165 |
| TpC ratio | 1.055 | 0.965 | 0.946 | 0.955 | 0.984 |
| CpA ratio | 1.068 | 1.106 | 1.118 | 1.113 | 1.096 |
| CpG ratio | 1.005 | 0.916 | 0.950 | 0.896 | 0.907 |
| CpT ratio | 1.013 | 0.944 | 0.929 | 0.948 | 0.953 |
| CpC ratio | 0.893 | 0.957 | 0.913 | 0.960 | 0.993 |
| MFE | -32.200 | -19.700 | -21.100 | -19.100 | -19.000 |
| MFED | 0.032 | 0.005 | 0.008 | 0.004 | 0.004 |

**Table S1. Distributions of Composition Metrics.**
| Metric | B) Comparison |  |  |  |  |
| --- | --- | --- | --- | --- | --- |
|  | Natural vs Transformer | Natural vs High quality | High quality vs Medium quality | High quality vs Low quality | Medium quality vs Low quality |
| GC Content | 5.93E-146 | 8.94E-32 | 1.77E-06 | 1.53E-11 | 0.06 |
| ApA ratio | 3.75E-04 | 0.67 | 0.21 | 1.84E-06 | 8.88E-04 |
| APG ratio | 3.32E-15 | 3.40E-04 | 0.02 | 0.23 | 0.23 |
| ApT ratio | 1.12E-10 | 4.90E-10 | 1.83E-05 | 3.99E-09 | 0.03 |
| ApC ratio | 8.57E-188 | 3.23E-52 | 4.11E-03 | 3.52E-09 | 3.74E-03 |
| GpA ratio | 3.17E-30 | 1.97E-09 | 0.58 | 0.03 | 0.14 |
| GpG ratio | 5.57E-38 | 1.45E-04 | 5.07E-04 | 4.39E-13 | 5.40E-05 |
| GpT ratio | 3.55E-294 | 4.99E-117 | 4.80E-04 | 4.70E-10 | 2.24E-03 |
| GpC ratio | 7.99E-212 | 4.66E-81 | 0.96 | 0.04 | 0.04 |
| TpA ratio | 6.22E-31 | 1.23E-06 | 1.82E-03 | 1.16E-05 | 0.31 |
| TpG ratio | 0.07 | 2.12E-04 | 9.74E-03 | 4.91E-09 | 3.80E-03 |
| TpT ratio | 3.65E-217 | 8.14E-92 | 5.81E-05 | 1.37E-16 | 1.46E-05 |
| TpC ratio | 2.42E-94 | 3.73E-46 | 0.35 | 8.97E-06 | 1.83E-03 |
| CpA ratio | 1.05E-46 | 6.21E-22 | 0.82 | 9.88E-04 | 4.48E-04 |
| CpG ratio | 1.04E-44 | 2.12E-08 | 2.38E-03 | 8.67E-04 | 0.94 |
| CpT ratio | 2.81E-30 | 2.65E-15 | 0.04 | 4.43E-04 | 0.23 |
| CpC ratio | 1.73E-62 | 0.01 | 5.00E-08 | 1.72E-23 | 3.16E-06 |
| MFE | 1.06E-229 | 1.26E-62 | 3.41E-06 | 1.01E-08 | 0.42 |
| MFED | <2.2e-308 | 1.13E-116 | 3.77E-06 | 3.00E-06 | 0.83 |

**Figure S2.**
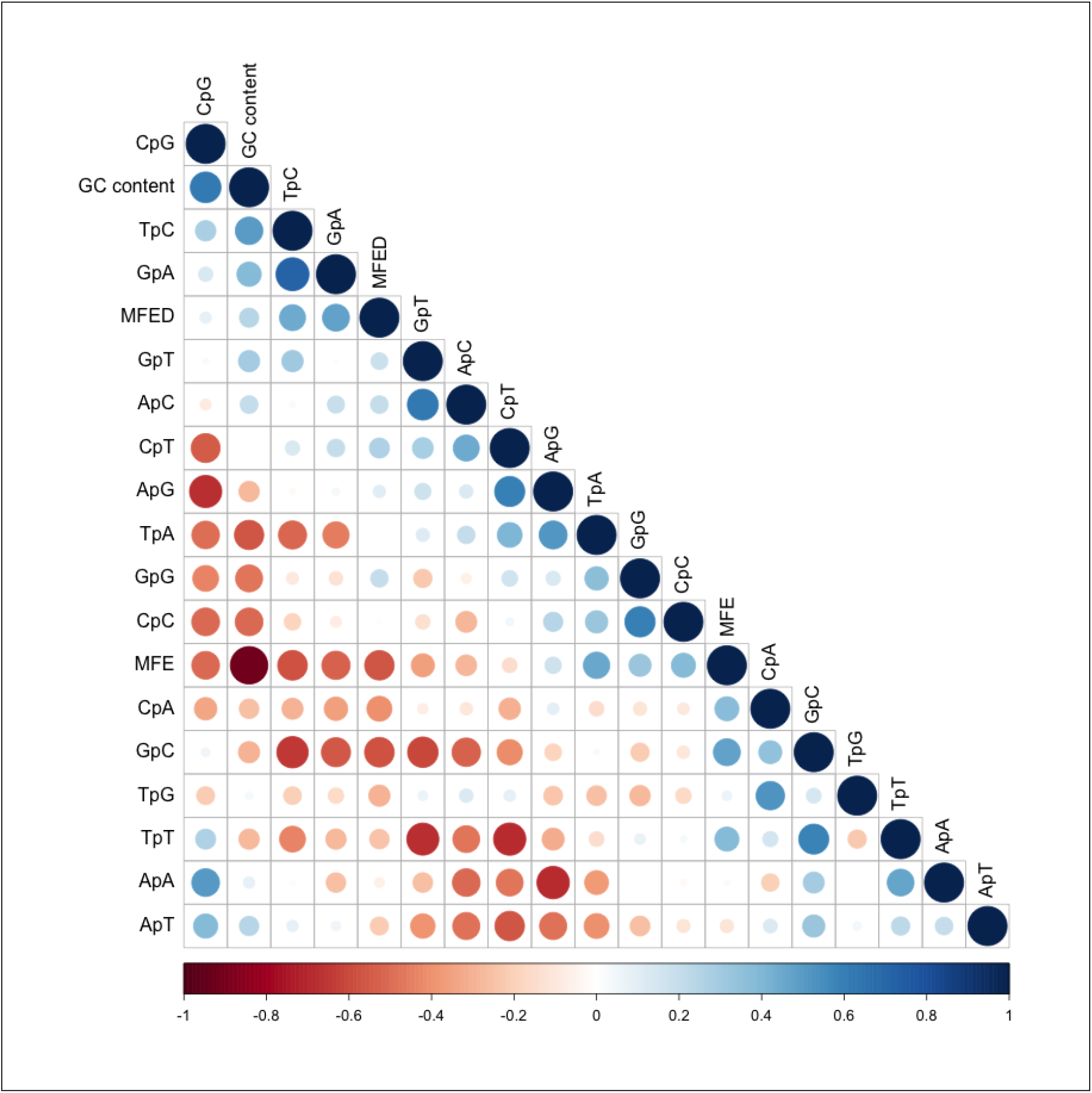
Correlation between Scaled Compositional Metrics. Correlations are inclusive of all data points regardless of completeness of taxonomy metadata. Correlation values calculated using function cor() from base R.

**Table S2.** Family-specific Clustering within PCA Results. “Nearest Neighbor Matches” defined as the number of family members for which the nearest data point in the 19-dimensional space (by weighted Euclidean distance) was a member of the same family. P value is derived from a binomial distribution where the expectation of matches is equal to the frequency of the family within the broader sample population (“Frequency Expected”). Note, this method is vulnerable to within-family heterogeneity, as a family with many small local clusters but large global variance would appear highly clustered with this method.

| Family | Nearest Neighbor Matches | Total Members | Frequency Observed | Frequency Expected | P Value |
| --- | --- | --- | --- | --- | --- |
| Ackermannviridae | 46 | 50 | 0.92 | 0.02 | 1.41E-80 |
| Aliceevansviridae | 93 | 93 | 1 | 0.03 | <2.2e-308 |
| Autographiviridae | 343 | 352 | 0.97 | 0.11 | <2.2e-308 |
| Blumeviridae | 5 | 34 | 0.15 | 0.01 | 1.86E-06 |
| Casjensviridae | 44 | 53 | 0.83 | 0.02 | 3.05E-71 |
| Chaseviridae | 29 | 30 | 0.97 | 0.01 | 4.45E-61 |
| Cystoviridae | 14 | 21 | 0.67 | 0.01 | 1.65E-28 |
| Demerecviridae | 84 | 84 | 1 | 0.03 | <2.2e-308 |
| Drexlerviridae | 105 | 108 | 0.97 | 0.04 | 2.87E-151 |
| Herelleviridae | 94 | 95 | 0.99 | 0.03 | 2.78E-144 |
| Inoviridae | 47 | 55 | 0.85 | 0.02 | 2.12E-76 |
| Kyanoviridae | 35 | 37 | 0.95 | 0.01 | 2.63E-68 |
| Mesyanzhinoviridae | 13 | 14 | 0.93 | 0 | 1.59E-33 |
| Microviridae | 25 | 30 | 0.83 | 0.01 | 1.31E-48 |
| Orlajensenviridae | 11 | 11 | 1 | 0 | <2.2e-308 |
| Peduoviridae | 84 | 92 | 0.91 | 0.03 | 1.67E-120 |
| Rountreeviridae | 36 | 38 | 0.95 | 0.01 | 8.70E-70 |
| Salasmaviridae | 26 | 29 | 0.9 | 0.01 | 7.71E-53 |
| Schitoviridae | 89 | 94 | 0.95 | 0.03 | 1.04E-130 |
| Steigviridae | 13 | 14 | 0.93 | 0 | 1.59E-33 |
| Steitzviridae | 403 | 412 | 0.98 | 0.13 | <2.2e-308 |
| Straboviridae | 151 | 154 | 0.98 | 0.05 | 1.69E-194 |
| Suoliviridae | 33 | 36 | 0.92 | 0.01 | 1.21E-63 |
| Tectiviridae | 7 | 10 | 0.7 | 0 | 5.50E-19 |
| Vilmaviridae | 29 | 29 | 1 | 0.01 | <2.2e-308 |
| Zierdtviridae | 19 | 19 | 1 | 0.01 | <2.2e-308 |
| Zobellviridae | 11 | 12 | 0.92 | 0 | 1.21E-29 |

**Figure S3.**
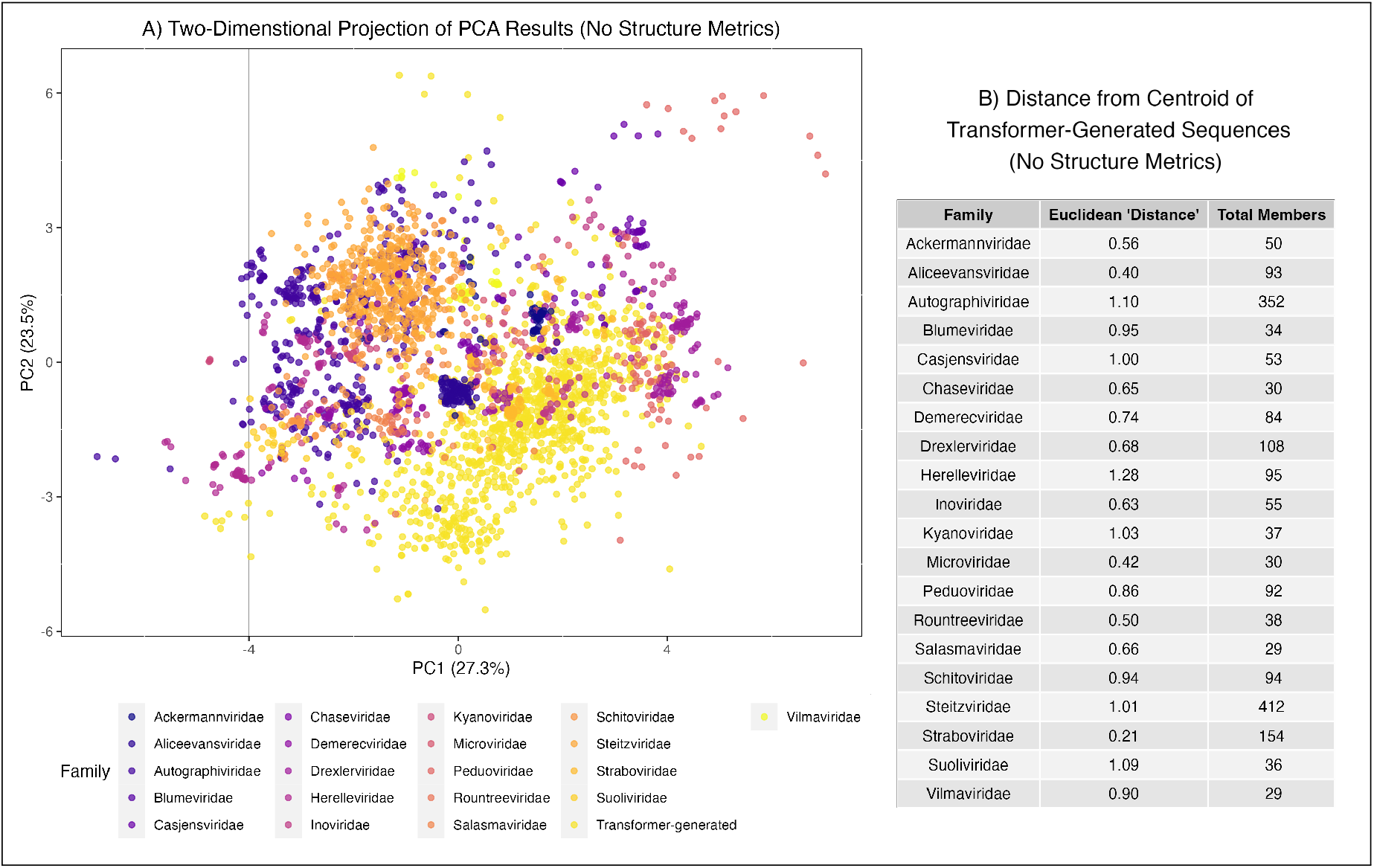
Principal Component Analysis (PCA) of Compositional Metrics without MFE or MFED. (**A**) Two-dimensional projections of PCs 1 and 2 limited to sequences from families with *≥* 25 members, colored by taxonomy. Percentages in x-axis and y-axis titles indicate percentage of variance explaned by given PC. Kindly note that the two-dimensional projection can be misleading as to the true location of a data point in the 19-dimensional space created by the PCA. Transformer-generated sequences clustered at a rate of 92.7% (929/1002). (**B**) Unitless weighted euclidean distance measures from centroid of family to centroid of transformer-generated sequences.

**Figure S4.**
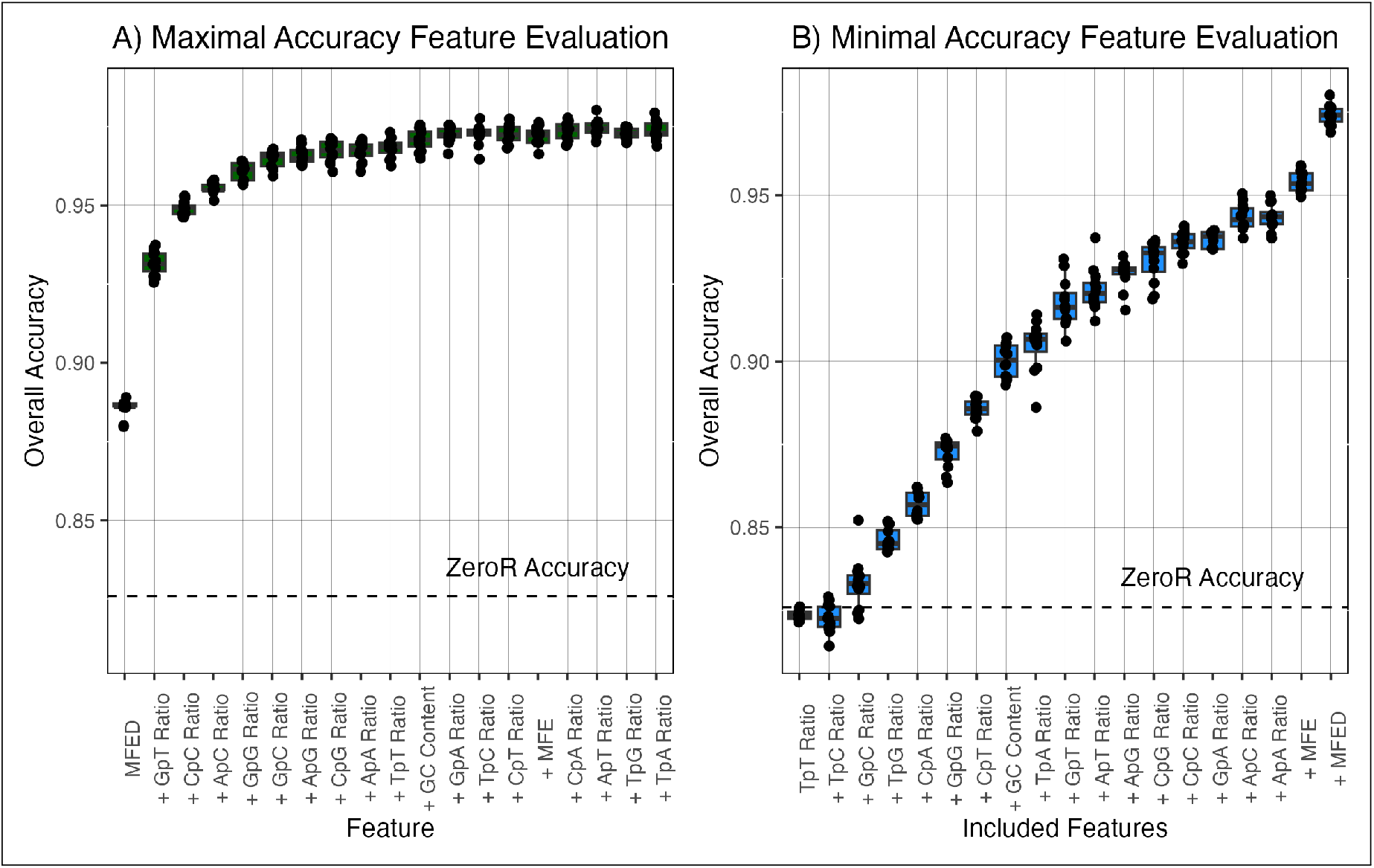
Neural Network Predictive Feature Evaluation. Results of predictive feature evaluation for (**A**) maximal and (**B**) minimal feature orders. X-axis labels specify the order in which features were added, with each point on the x-axis specifying the newest feature. Each model was inclusive of all features preceding that point on the x-asix.

**Figure S5.**
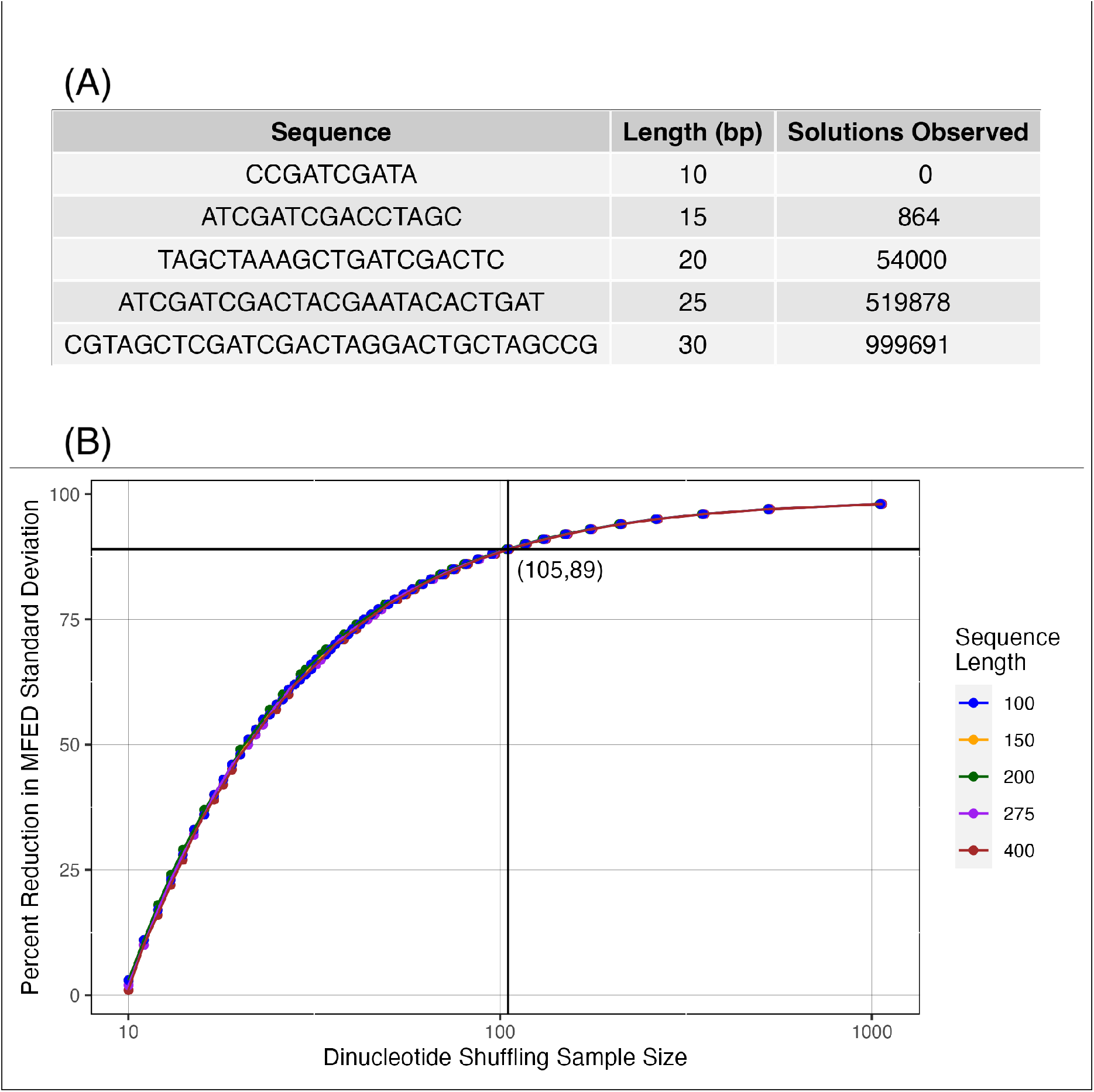
Dinucleotide Shuffling Sample Size Optimization. (**A**) Results of dinucleotide shuffling observed for each random input sequence. Solutions observed are number of unique dinucleotide shuffled sequence outputs after performing n = 1,000,000 independent shuffles. (**B**) Relationship between shuffling sample size and variability of MFED estimate. Values obtained as described in Supplementary Methods. Vertex displays sample size implemented for this study.

**Figure S6.**
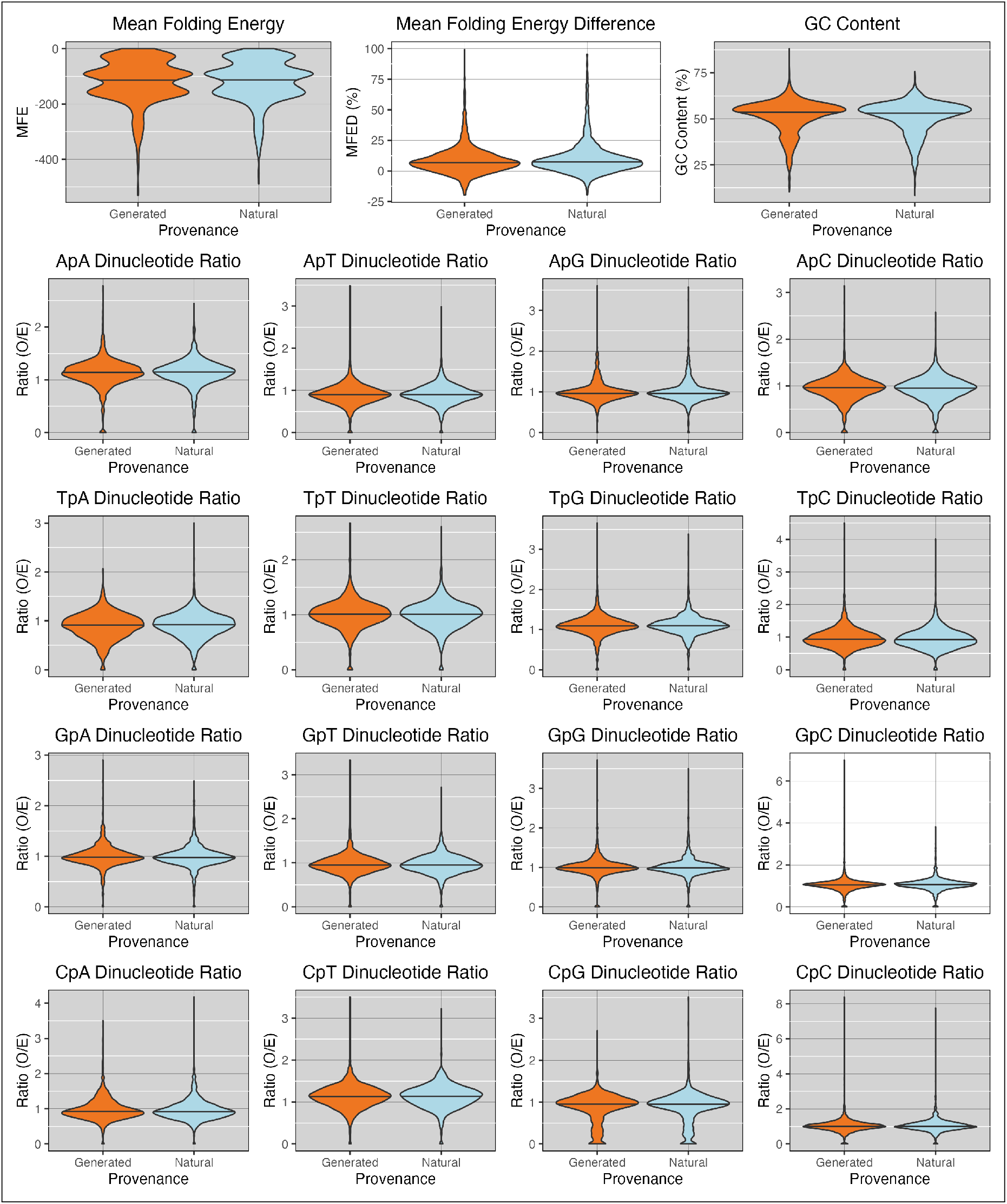
Comparisons of Compositional Metrics between Natural and GenerRNA Sequences. Only MFED (p = 0.00062) and GpC ratio (p = 0.0017) had significantly different distributions by two-tailed Mann-Whitney U Test after a Bonferroni correction. For ease of viewing, these panels are colored white.

**Figure S7.**
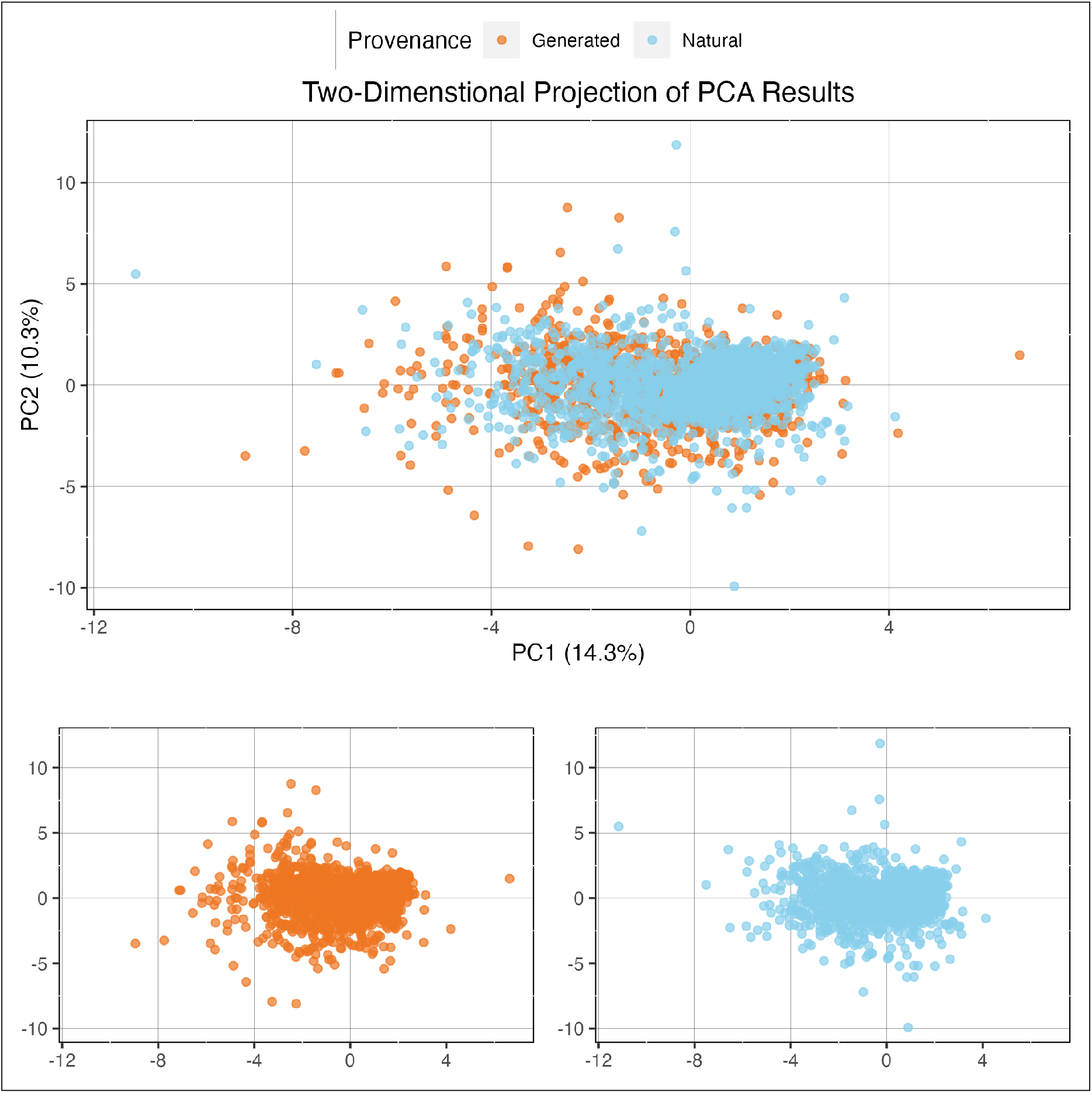
Principal Component Analysis (PCA) of Compositional Metrics for GenerRNA sequences. Two-dimensional projection of PCA results for all natural and generated sequences in the GenerRNA dataset. Views of “generated only” and “natural only” provided due to substantial overlap in the main panel. Neither generated nor natural sequences clustered within this analysis by binomial distribution tests.

## Notes

### Competing Interest Statement

The authors have declared no competing interest.

